# Improved detection of genetic effects on promoter usage with augmented transcript annotations

**DOI:** 10.1101/2022.07.12.499800

**Authors:** Andreas Vija, Kaur Alasoo

**Affiliations:** Institute of Computer Science, University of Tartu, Tartu, Estonia; STACC OÜ, Tartu, Estonia

## Abstract

Disease-associated non-coding variants can modulate their target genes by disrupting multiple mechanisms, including regulating total gene expression level, splicing, alternative polyadenylation or promoter usage. Quantifying promoter usage from standard RNA sequencing data is challenging due to incomplete reference transcriptome annotations and low read coverage observed at the ends of transcripts. We previously developed the txrevise tool (https://github.com/kauralasoo/txrevise) to quantify promoter usage events from RNA-seq data using reference transcriptome annotations. Here, we augment the txrevise promoter event annotations with experimentally identified Cap Analysis of Gene Expression (CAGE) promoters from the FANTOM5 project. Applying the new annotations to RNA-seq data from 358 individuals, we found that augmented promoter event annotations increased the power to detect promoter usage quantitative trait loci (puQTLs) by ~30%. However, concordance between puQTLs inferred from RNA-seq data and those directly measured using CAGE remained low, suggesting that additional experimental and computational improvements are needed to capture the full range of regulatory effects of non-coding variants.

## Introduction

Genetic variants regulating promoter usage can play an important role in human complex traits (Alasoo et al., 2019; Garieri et al., 2017; Kubota and Suyama, 2022). Promoter usage can be directly quantified using experimental techniques that capture 5’ ends of transcripts such as Cap Analysis of Gene Expression (CAGE) (Shiraki et al., 2003), but currently only one such population-level human dataset exists (Garieri et al., 2017). Alternatively, promoter usage can be quantified from standard bulk RNA sequencing (RNA-seq) data (Figure 1a). The advantage of this approach is that bulk RNA-seq data is readily available from thousands of individuals and over a hundred different cell types or tissues (Kerimov et al., 2021; The GTEx Consortium, 2020).

**Figure 1.**
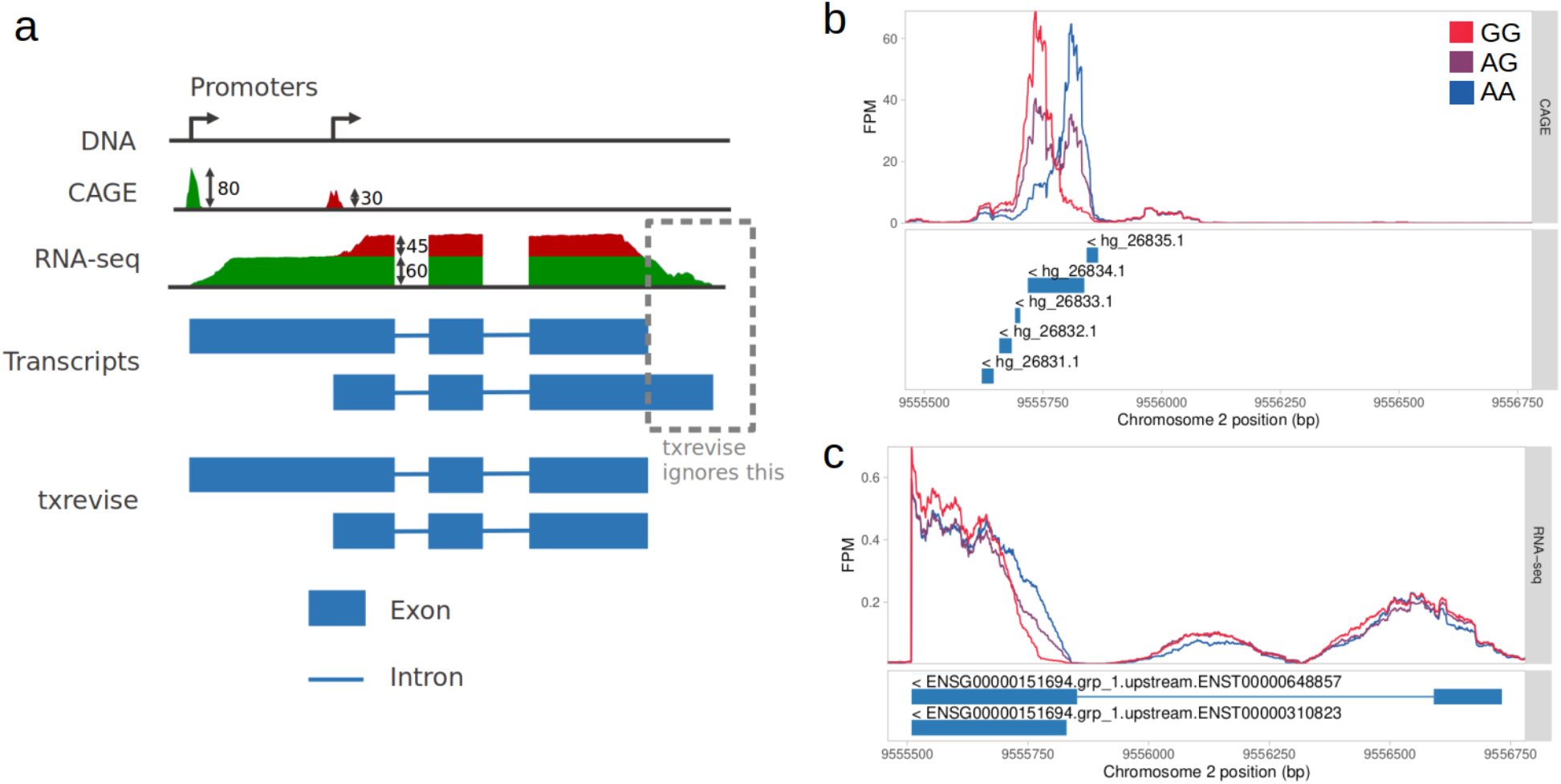
Comparison of RNA sequencing methods for promoter usage quantification. (**a**) Fictional gene with two alternative promoters. CAGE directly sequences the 5’ ends of expressed transcripts and thus detects two peaks corresponding to the two alternative promoters. Promoter usage is quantified by counting the reads overlapping the two peaks. RNA-seq captures reads across the full length of the expressed transcripts with a notable drop at the beginning. Most reads originating from this gene are compatible with both known transcripts. Promoter usage can be quantified by estimating the relative expression of the two transcripts that best explains the observed read coverage pattern across the gene. However, transcript annotations often couple alternative promoters with unrelated splicing or 3’ end events. Txrevise overcomes this by constructing new annotations corresponding to independent promoter usage events. (**b**) Promoter usage QTL affecting *ADAM17* (ENSG00000151694). The CAGE read coverage stratified by the genotype of the lead puQTL variant (rs12692386) shows strong genotype-dependent shift in promoter usage. (**c**) RNA-seq read coverage signal captures similar change, but this does not correspond to any annotated *ADAM17* promoters.

However, quantification of promoter usage from standard RNA-seq data is complicated by multiple factors. First, read coverage is much lower at the ends of transcripts (Love et al., 2016; Roberts et al., 2011), which makes it difficult to precisely detect the location of each transcription start site (TSS) (Pertea et al., 2015). Consequently, it is often hard to ascertain more than one TSS per gene from RNA-seq data (Adiconis et al., 2018). Secondly, transcripts contain overlapping exons which means that most reads cannot be uniquely assigned to a specific transcript (Figure 1a). Thus, methods for quantifying promoter usage from RNA-seq such as txrevise (Alasoo et al., 2019) and proActiv (Demircioğlu et al., 2019) rely heavily on pre-existing promoter annotations (Figure 1a) from databases such as Ensembl (Howe et al., 2021). This means that even if a genetic effect on promoter usage is clearly visible from RNA-seq read coverage track, it might not be detected by existing methods due to incomplete promoter annotations (Figure 1c).

Previous research has demonstrated that detection of alternative polyadenylation events can be significantly improved by incorporating experimental annotations such as data from 3’ RNA-seq experiments (Ha et al., 2018; Shah et al., 2021). Here, we augment txrevise promoter annotations using experimental CAGE (Shiraki et al., 2003) data from the FANTOM5 project (FANTOM Consortium and the RIKEN PMI and CLST et al., 2014). We demonstrate that incorporating experimentally detected promoter annotations improves concordance between CAGE and RNA-seq data and increases the number of detected promoter usage quantitative trait loci (puQTLs) by around 30%. Nevertheless, overall concordance between puQTLs detected by CAGE and RNA-seq remains low, suggesting that the two approaches have distinct strengths and weaknesses.

## Results

### Concordance of puQTLs detected by CAGE and RNA-seq

To assess the concordance between the puQTLs detected by CAGE and RNA-seq, we re-analysed two transcriptomic datasets generated from lymphoblastoid cell lines. The Garieri_2017 (Garieri et al., 2017) dataset contained CAGE data from 154 individuals from the 1000 Genomes (1000 Genomes Project Consortium et al., 2015) and GENCORD (Gutierrez-Arcelus et al., 2013) cohorts. The GEUVADIS (Lappalainen et al., 2013) datasets contained RNA-seq data from 358 individuals from the 1000 Genomes cohort. All individuals were of European ancestries and 78 individuals were shared between the two datasets.

We first compared CAGE against RNA-seq using Ensembl reference promoter annotations. Alternative promoter annotations were extracted from reference transcriptome with txrevise (Alasoo et al., 2019). We found that using the same +/− 200 kb *cis* window and 5% false discovery rate (FDR) sigificance threshold, CAGE detected at least one significant puQTL for more genes than txrevise (1145 vs 979 genes), even though the RNA-seq dataset was more than two times larger (154 vs 358 samples) (Table 1). When varying the gene expression threshold, we found that CAGE was more sensitive for lowly expressed genes whereas txrevise found more associations at higher gene expression thresholds (Table 1, Supplementary Figure 1). Nevertheless, the agreement between the two datasets was small with only 307 genes having a significant puQTL in both datasets (Figure 2a). An example puQTL for *ADAM17* detected by CAGE and missed by txrevise is illustrated on Figure 2b. Close inspection of the corresponding read coverage plots (Figure 1b-c) revealed that while the puQTL signal was clearly visible from the RNA-seq data, the association was missed because neither of the two annotated promoters (corresponding to transcripts ENST00000648857 and ENST00000310823) overlapped the downstream alternative promoter captured by CAGE.

**Table 1.** Number of puQTLs (FDR < 0.05) detected at various gene expression thresholds by CAGE, reference-based txrevise and augmented txrevise. As transcripts per million (TPM) > 1 allowed filtering out many lowly expressed genes without a major drop in the number of puQTLs detected, it was chosen as the threshold.

|  |  | Number of puQTLs detected |  |  |  |
| --- | --- | --- | --- | --- | --- |
| Min TPM threshold | % of genes above threshold | CAGE | txrevise (reference only) | txrevise (augmented) | puQTL % increase |
| None | 100% | 1208 | 1073 | 1393 | 29.8% |
| 0.1 | 79.4% | 1235 | 1051 | 1359 | 29.3% |
| 1 | 62.9% | 1145 | 979 | 1287 | 31.5% |
| 10 | 31.5% | 564 | 663 | 868 | 30.9% |
| 100 | 4.8% | 87 | 152 | 201 | 32.2% |

**Figure 2.**
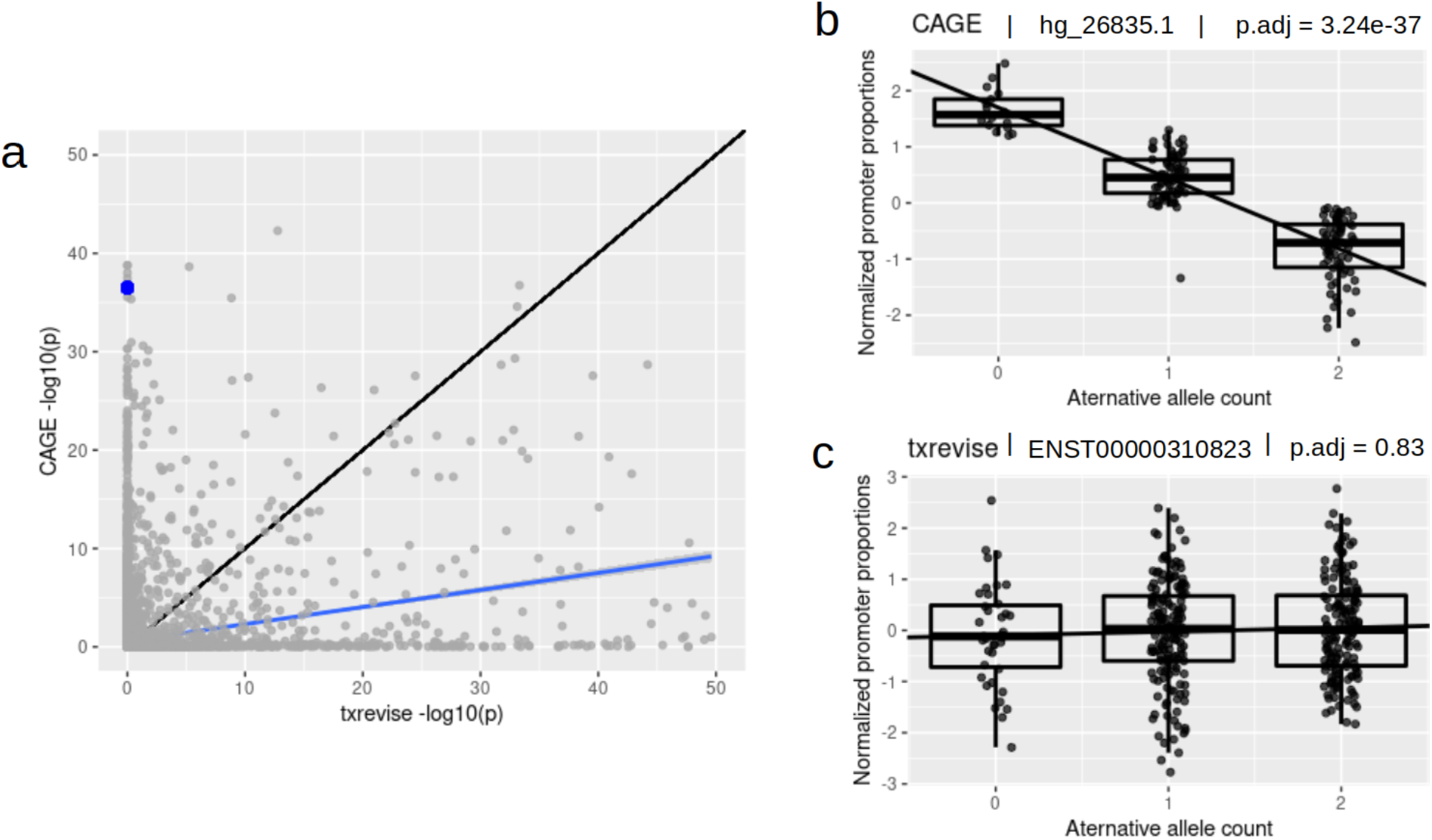
Concordance of promoter usage QTLs detected by CAGE and txrevise. (**a**) Scatterplot of the lead variant p-values of each gene from CAGE and reference-based txrevise analysis. The identity line (black) diverges from the regression line (blue), indicating low concordance between the two methods. The blue dot corresponds to the lead puQTL variant of *ADAM17*, rs12692386. The variant was significantly associated with *ADAM17* promoter usage in CAGE analysis but not in txrevise analysis. (**b**) Normalised usage of the *ADAM17* CAGE promoter hg_26835.1 stratified by the genotype of the puQTL lead variant (rs12692386). (**c**) Normalised usage of the *ADAM17* ENST0000031082 txrevise reference promoter stratified by the genotype of the puQTL lead variant (rs12692386).

### Augmenting promoter annotations using CAGE data

To overcome the limitation of incomplete promoter annotations in txrevise analysis, we obtained a list of experimentally detected promoters from the FANTOM5 project (Abugessaisa et al., 2017) and used a simple heuristic approach to construct novel transcript annotations based on these promoters (Figure 3) (see Methods). Briefly, we used the FANTOM5 promoters to construct new alternative first exons if the novel promoters were in an existing exon or 1000 bp upstream of one and if the newly constructed promoter was at least 20 nucleotides away from any existing promoter. This process increased the number of txrevise alternative promoter annotations from 72,292 to 114,768 (37%). Furthermore, 1650 genes that previously had only one annotated promoter now had more than one, thus enabling promoter usage quantification for those genes.

**Figure 3.**
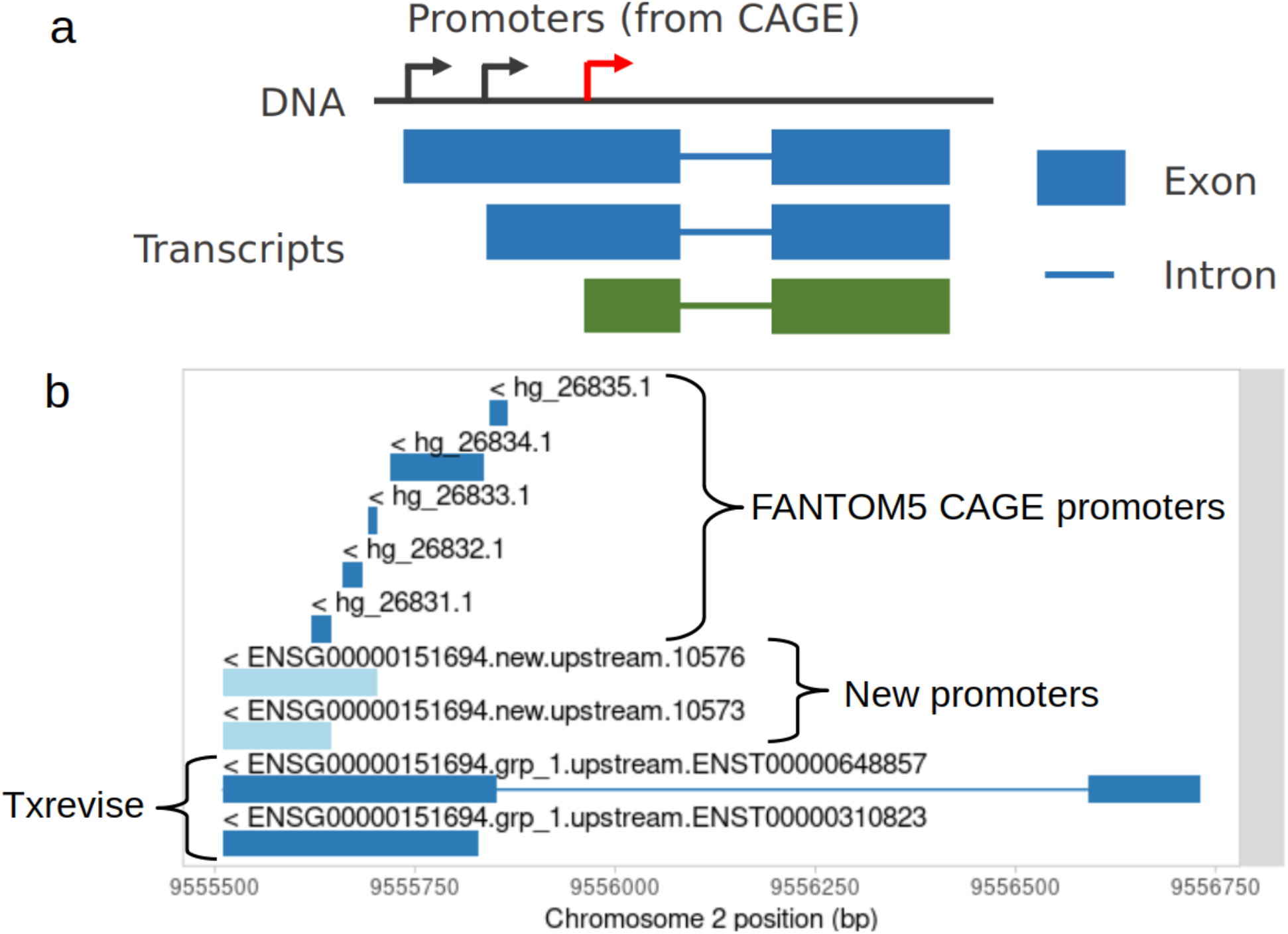
Constructing new transcript annotations based on CAGE peaks. (**a**) Fictional gene with two alternative promoters (black) corresponding to two transcripts (blue) starting from those promoters, but also an additional promoter (red) that has no existing transcript annotation but for which a hypothetical transcript (green) could be constructed based on existing transcripts. (**b**). Two new txrevise promoter annotations (light blue) constructed for *ADAM17* by augmenting existing txrevise promoter annotations (dark blue, bottom) with CAGE promoters (dark blue, top) from FANTOM5.

### Impact of augmented promoter annotations on puQTL detection

Next, we re-analysed the GEUVADIS dataset using the augmented txrevise promoter annotations. We found that augmented annotations increased puQTL yield by ~30% at all gene expression level thresholds (Table 1, Figure 4a). Similarly, the number of shared puQTLs genes detected by both CAGE and txrevise increased from 307 to 397 (Figure 4b). One such example was the previously missed *ADAM17* gene, but even with augmented annotations, the association detected from RNA-seq data (Figure 4c-d) was much weaker compared to the CAGE signal (Figure 2b). Similarly, the overall concordance between CAGE and txrevise still remained relatively low (Figure 4b).

**Figure 4.**
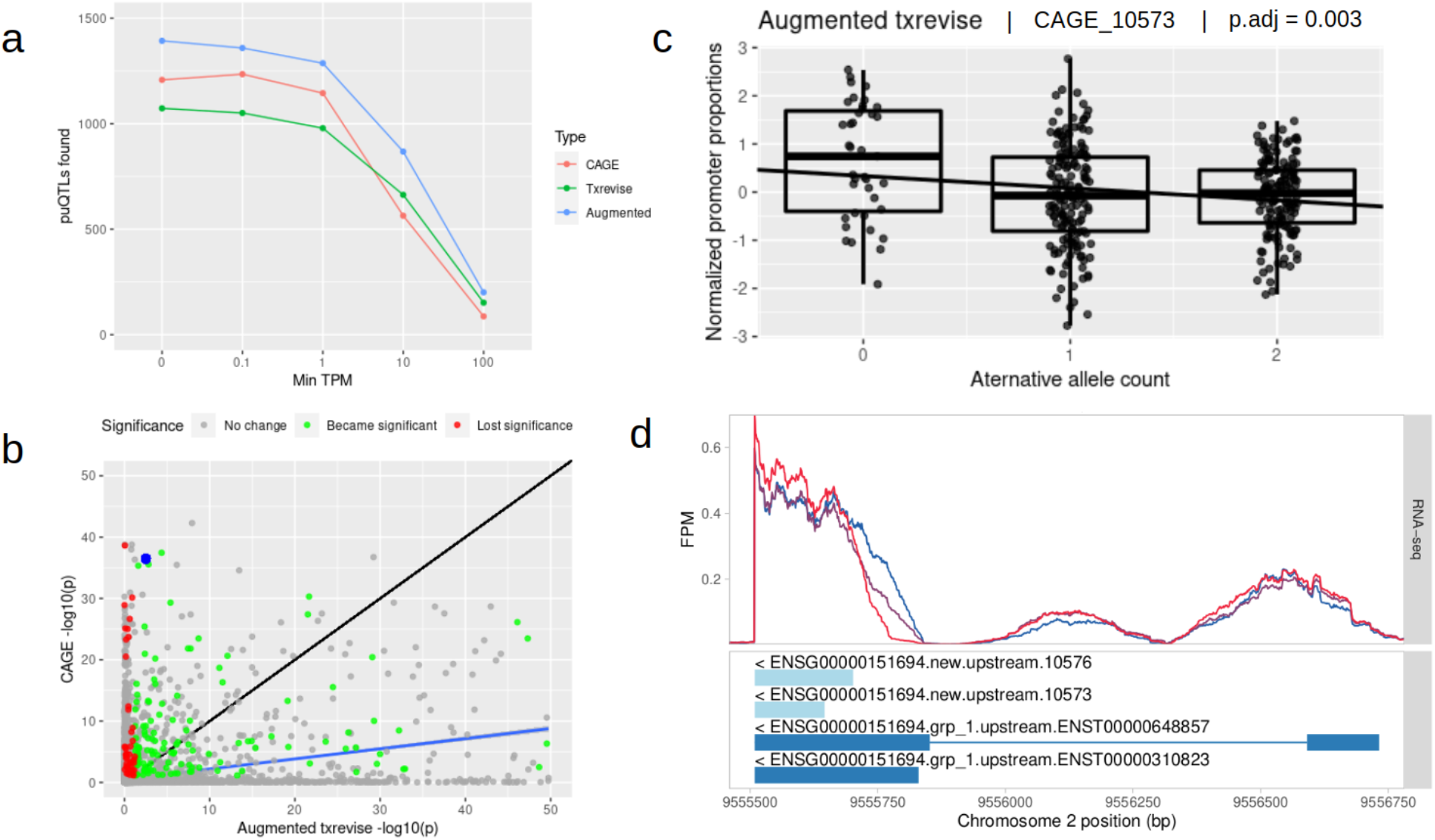
Impact of augmented promoter annotations on puQTL detection. (**a**) Number of genes with at least one significant puQTL (y-axis) as a function of gene expression level threshold (x-axis) (TPM - transcripts per million). (**b**) The lead variant p-values of each gene for CAGE and txrevise with an identity line (black) and a regression line (blue). Green dots represent genes for which a significant puQTL was detected only after promoter augmentation, red dots represent genes that lost a significant puQTL after augmentation. The blue dot corresponds to the lead variant of *ADAM17*, rs12692386. (**c**) Normalised usage of the newly added ENSG00000151694.new.upstream.10573 (CAGE_10573) txrevise promoter stratified by the genotype of the puQTL lead variant (rs12692386) (**d**) RNA-seq read coverage at the *ADAM17* promoter stratified by the genotype of the puQTL lead variant (rs12692386). The GG genotype (red line) is associated with a shift towards an upstream promoter relative to the AA genotype (blue line). The newly added promoter annotation (ENSG00000151694.new.upstream.10573, light blue) can better capture this shift compared to the two existing reference-based promoters (dark blue).

## Discussion

We performed puQTL analysis in lymphoblastoid cell lines using two complementary technologies: CAGE that directly sequences 5’ ends of transcripts and txrevise that can leverage reference transcript annotations to capture promoter usage events from fulllength RNA-seq data. We found that the concordance in the puQTLs detected with the two approaches was generally low. Augmenting reference transcript annotations with novel FANTOM5 promoters increased the ability of txrevise to detect puQTLs by 30%, but concordance with CAGE puQTLs still remained low. We believe that the discordance is primarily due to differences in technology with CAGE being able to better distinguish promoters of lowly expressed genes and having higher signal-to-noise ratio due to more direct measurement. However, since the CAGE and RNA-seq data were generated by two independent studies, we cannot rule out that other technical factors might contribute to the observed differences.

Our results indicate that a significant proportion of alternative promoter annotations are still missing from the Ensembl database. Consequently, we found that augmenting reference transcripts with experimentally determined promoters from the FANTOM5 project significantly increased the number of puQTLs detectable from RNA-seq data. As a result, using augmented promoter annotation to re-process publicly available RNA-seq eQTL datasets by projects such as the eQTL Catalogue (Kerimov et al., 2021) has a great potential to increase the number of puQTLs detected. Future studies can explore if incorporating additional experimentally derived promoter annotations such as those generated by the RAMPAGE project (Moore et al., 2021) or transcriptome assembly methods (Kubota and Suyama, 2022) can further improve the ability to detect puQTLs from RNA-seq data.

## Methods

### FANTOM5 promoter annotations

We downloaded annotations of 210,250 human promoters from the FANTOM5 database (Abugessaisa et al., 2017; FANTOM Consortium and the RIKEN PMI and CLST et al., 2014), which was constructed based on CAGE peaks. Of these, 96,562 promoter annotations associated with an autosomal gene were kept. The name of the associated gene was mapped to an Ensembl id using the eQTL Catalogue gene metadata files (https://doi.org/10.5281/zenodo.3366011). Of the remaining annotations, 93,663 promoters from 20,201 genes had a gene name which mapped to exactly one Ensembl id. Of these, 93,554 promoters from 20,193 genes were mapped to a unique chromosome and were thus used in further analysis.

### GAGE data processing

We downloaded the raw CAGE sequencing data from the Garieri_2017 (Garieri et al., 2017) study from ArrayExpress (E-MTAB-5835). CAGE reads were mapped to the GRCh38 human reference genome using Burrows-Wheeler Aligner v0.7.12 (Li and Durbin, 2009) and multi-mapping reads were discarded. The number of CAGE reads corresponding to each promoter was counted using featureCounts (Liao et al., 2014). On average, 46.0% of all CAGE reads overlapped with the TSS of some FANTOM5 promoter.

Based on these read counts, we omitted from further analysis all CAGE promoters with zero total mapped reads across all samples. After this, 90,003 promoters from 18,546 genes remained.

To quantify promoter usage and not general gene expression in CAGE data, the number of reads assigned to each promoter was divided by the total number of reads assigned to all promoters of the same gene. Missing promoter usage values were replaced by the mean calculated across all individuals. Finally, rank-based inverse normal transformation was used to enable more robust use of linear models (McCaw et al., 2019).

### Genotype data processing

We re-analyzed genotype data from 154 individuals of European descent from the Garieri_2017 study, 86 of which were from the 1000 Genomes Project (1000 Genomes Project Consortium et al., 2015) and 68 from the GENCORD project (Gutierrez-Arcelus et al., 2013). Genotypes from the GENCORD project were imputed to the 1000 Genomes reference panel as described previously (Kerimov et al., 2021). Whole genome sequencing genotypes for the 1000 Genomes Project samples were downloaded from the 1000 Genomes Project FTP server (http://ftp.1000genomes.ebi.ac.uk/vol1/ftp/). The 9.2 million genetic variants shared by these two datasets were used in the rest of the analysis.

### RNA-seq data processing

Txrevise promoter usage events (both original and augmented) were quantified with the <u>eQTL-Catalogue/rnaseq</u> workflow and normalised with the <u>eQTL-Catalogue/qcnorm</u> workflow as described previously (Kerimov et al., 2021). Only the 15,275 genes present in both txrevise transcript annotations and the filtered FANTOM5 annotations were used for downstream analysis.

Genes with very low expression levels are likely to be biologically insignificant and their expression estimates are vulnerable to noise. Gene expression levels were calculated based on the output of featureCounts generated during the execution of the <u>eQTL-Catalogue/rnaseq</u> pipeline on the RNA-seq data. For various TPM values, only genes with at least 5% of individuals exhibiting at least that much expression were considered.

### Promoter usage QTL analysis

To map puQTLs, we used the <u>eQTL-Cataloge/qtlmap</u> Nextflow workflow built on top of fastQTL (Ongen et al., 2016) and QTLtools (Delaneau et al., 2017) by the eQTL Catalogue project (Kerimov et al., 2021). For every gene, only genetic variants within the +/− 200kb *cis* window centred around the canonical Ensembl promoter of the gene were considered. The coordinates of genes were obtained from the eQTL Catalogue gene metadata files (https://doi.org/10.5281/zenodo.3366011). Multiple testing correction was performed as described previously (Kerimov et al., 2021).

### Creating new transcript annotations

All FANTOM5 promoters meeting the following criteria for different values of N were chosen:

- The promoter is not within N bp of the start of the exon of any existing txrevise transcript (taking into account the strand of the transcript)
- The promoter is not within N bp of any already added promoter
- The promoter overlaps with an exon of an existing txrevise transcript or is within 1000 bp upstream of one (94.4% of all promoters corresponding to a gene possessing a txrevise transcript match this criterion)

The chosen promoters and the txrevise transcripts whose exon the promoters overlapped or were near upstream to were used to construct a new set of artificial transcript annotations. The first exon of these new transcripts was an artificial exon from the promoter’s start coordinate to the end coordinate of the nearest (overlapping) first exon and the remaining exons were all the remaining exons of the existing txrevise transcript (Figure 3).

The created transcripts were added to txrevise transcripts, given as input to another run of txrevise and put through the <u>eQTL-Cataloge/qtlmap</u> workflow. Supplementary Figure 2 shows that the smaller the N gets, the more statistically significant puQTL genes were found, especially at values of N of 20 and smaller. For values of N larger than 100, the effect of adding annotations was negative, likely because re-running txrevise with N values larger than 25 causes some original txrevise transcripts to be removed. The biggest fraction of new genes that were also significant with CAGE occurred at N values of 5-25 and the least originally found genes were lost when N was 25. For these reasons, we chose 20 as the optimal N value based on our tests. This resulted in 77,869 new transcripts across 11,877 genes.

## Supplementary Figures

**Supplementary Figure 1.**
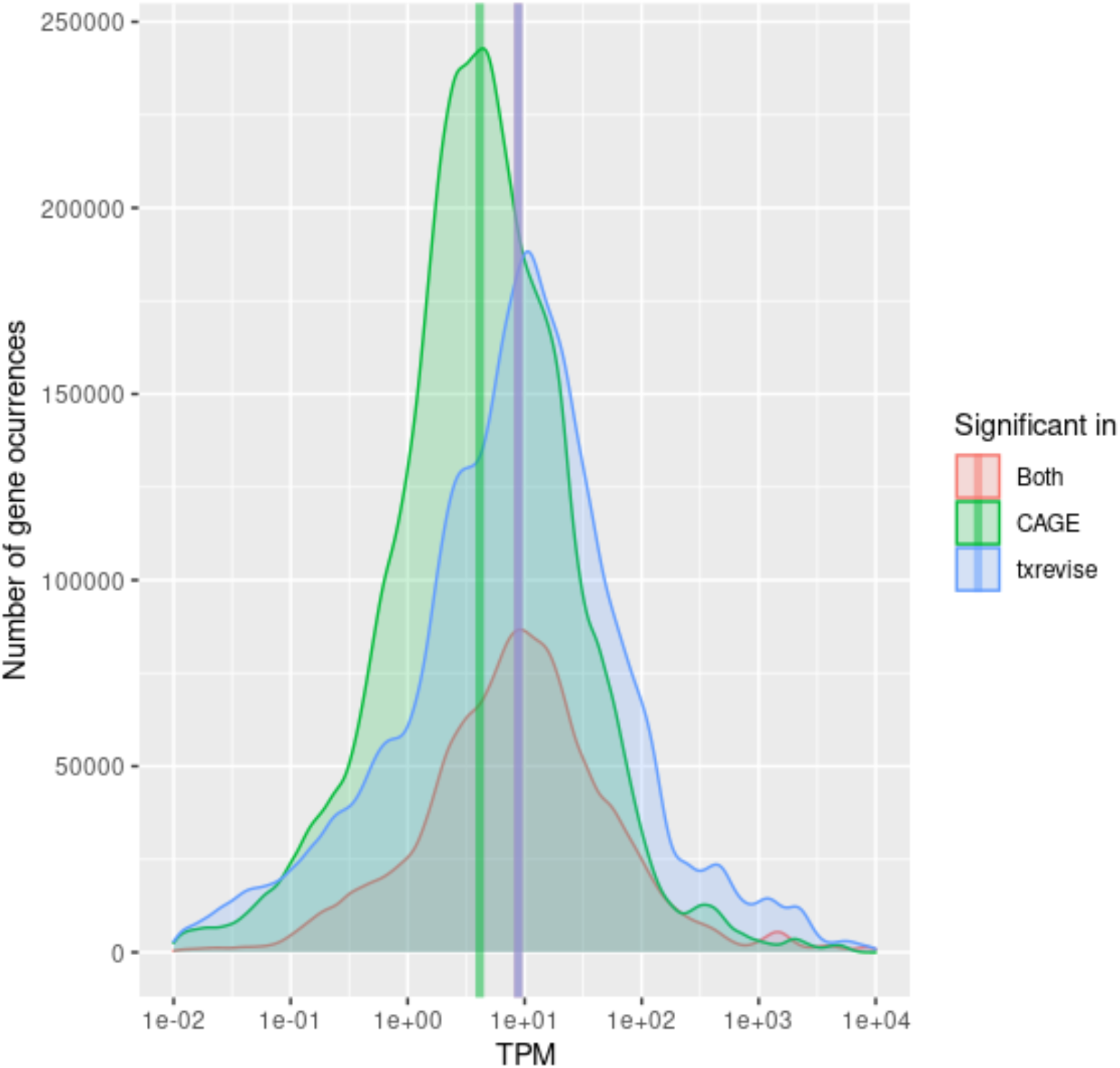
The distribution of transcripts per million (TPM) values across all significant puQTL genes grouped by whether the gene was found to be significant in CAGE or txrevise. Vertical lines show the median TPM of the corresponding group. The genes detected by CAGE have significantly lower mean TPM values than the genes detected by txrevise (Mann-Whitney U test p-value < 2.2e-16).

**Supplementary Figure 2.**
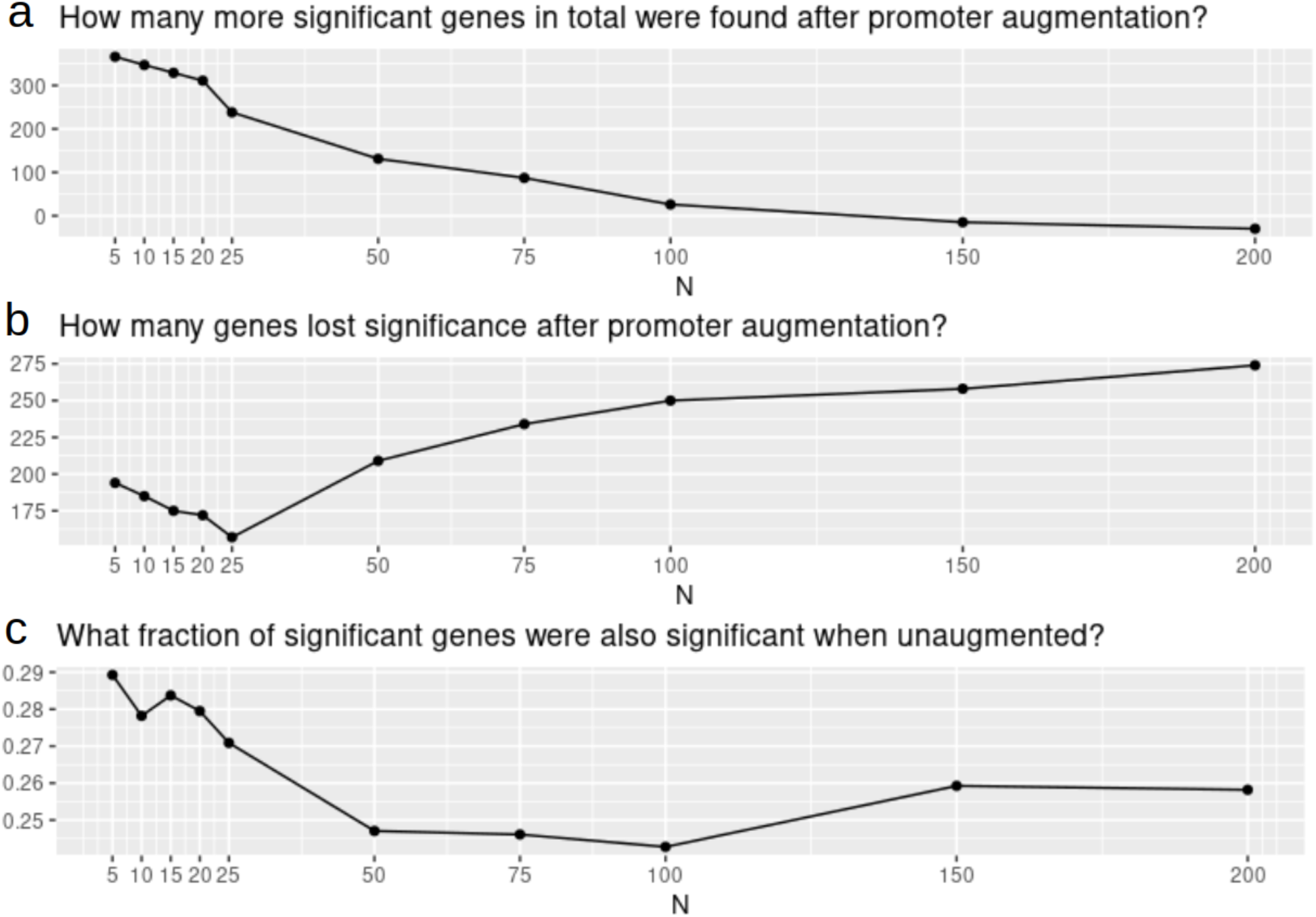
Effect of different values of N on detecting genes with at least one puQTL. (**a**) The effect of N on the number of additional puQTL genes detected. (**b**) The effect of N on the number of puQTL genes lost. (**c**) The effect of N on the agreement between reference and augmented and promoters.

## Data availability

The CAGE sequencing data from Garieri_2017 is available from ArrayExpress (E-MTAB-5835). The genotype data from the GENCORD study is available from EGA (EGAD00001000428). The RNA-seq data from the GEUVADIS study is available from ArrayExpress (E-GEUV-1). The GEUVADIS genotype data was downloaded from the 1000 Genomes Project FTP server (http://ftp.1000genomes.ebi.ac.uk/vol1/ftp/). The CAGE and txrevise promoter usage QTL summary statistics have been deposited to Zenodo (https://doi.org/10.5281/zenodo.5831090). The pre-computed txrevise annotations using Ensembl 105 transcriptome annotations and FANTOM5 promoters with N=25 parameter have been deposited to Zenodo (https://doi.org/10.5281/zenodo.6499127).

## Code availability

Source code of all the analyses presented in the paper is available from GitHub (https://github.com/andreasvija/cage). The updated version of the txrevise software supporting augmenting annotations with FANTOM5 CAGE promoters is available from GitHub (https://github.com/kauralasoo/txrevise). The txrevise promoter usage quantification was performed with the <u>eQTL-Catalogue/rnaseq</u> workflow and puQTL mapping was performed with the <u>eQTL-Catalogue/qtlmap</u> workflow.

## Acknowledgements

Many of the analyses were performed at the High Performance Computing Center, University of Tartu.

